# Joint population coding and temporal coherence link an attended talker’s voice and location features in naturalistic multi-talker scenes

**DOI:** 10.1101/2024.05.13.593814

**Authors:** Kiki van der Heijden, Prachi Patel, Stephan Bickel, Jose L. Herrero, Ashesh D. Mehta, Nima Mesgarani

## Abstract

Listeners effortlessly extract multidimensional auditory objects, such as a localized talker, from complex acoustic scenes. However, the neural mechanisms that enable simultaneous encoding and linking of distinct sound features—such as a talker’s voice and location—are not fully understood. Using invasive intracranial recordings in neurosurgical patients, we investigated how the human auditory cortex processes and integrates these features during naturalistic multi-talker scenes. We found that cortical sites exhibit a gradient of feature sensitivity, ranging from single-feature sensitive sites (responsive primarily to voice or location) to dual-feature sensitive sites (responsive to both features). At the population level, neural response patterns from both single- and dual-feature sensitive sites jointly encoded the attended talker’s voice and location. Notably, single-feature sensitive sites encoded their primary feature with greater precision but also represented coarse information about the secondary feature. Sites selectively tracking a single, attended speech stream concurrently encoded both voice and location features, demonstrating a link between selective attention and feature integration. Additionally, attention selectively enhanced temporal coherence between voice- and location-sensitive sites, suggesting that temporal synchronization serves as a mechanism for linking these features. Our findings highlight two complementary neural mechanisms—joint population coding and temporal coherence— that enable the integration of voice and location features in the auditory cortex. These results provide new insights into the distributed, multidimensional nature of auditory object formation during active listening in complex environments.

**SIGNIFICANCE STATEMENT:** In everyday life, listeners effortlessly extract individual sound sources from complex acoustic scenes which contain multiple sound sources. Yet, how the brain links the different features of a particular sound source to each other – such as a talker’s voice characteristics and location - is poorly understood. Here, we show that two neural mechanisms contribute to encoding and integrating voice and location features in multi-talker sound scenes: (1) some neuronal sites are sensitive to both voice and location and their activity patterns encode these features jointly; (2) the responses of neuronal sites that process only one sound feature – that is, location or voice – align temporally to form a stream that is segregated from the other talker.

**HIGHLIGHTS:**

- Auditory cortex exhibits a gradient of feature sensitivity, with some sites encoding only voice or location features, while others encode both simultaneously (dual-feature sensitive sites).
- Dual-feature sensitive sites integrate voice and location features of an attended talker with equal accuracy, providing a unified representation in multi-talker scenes.
- Single-feature sensitive sites primarily encode their preferred feature with high precision but also represent coarse information about other features, contributing to population-level integration.
- Temporal coherence selectively enhances synchronization between voice- and location-sensitive sites, providing another mechanism for integrating an auditory object’s features.
- Multi-dimensional auditory object formation relies on complementary neural mechanisms: joint population coding and temporal coherence.

## INTRODUCTION

In everyday life, listeners rapidly and effortlessly parse complex acoustic scenes with multiple sound sources into its individual constituents. This process of auditory scene analysis (ASA^1^) is based on the segregation and subsequent grouping of features of temporally overlapping sound sources, resulting in the formation of coherent auditory objects^2^. The sound features that contribute to such auditory object formation include voice features related to object identity (e.g., pitch or timbre) and location features (e.g., interaural time differences, location cues)^3–5^. However, the neural basis for multi-dimensional auditory object formation in complex, naturalistic listening scenes is poorly understood.

One unresolved question is where cortical representations of individual sound features are linked by the brain to form a multi-dimensional auditory object. In particular, as the prevailing dual-stream framework^6,7^ posits that voice and location features are encoded independently in two separate, functionally specialized and hierarchical processing streams it is not clear how these features are subsequently integrated to form a multi-dimensional auditory object. However, recent studies using an active task design indicate that sound feature encoding may be distributed across auditory cortex rather than taking place in dedicated, functionally specialized anatomical regions as posited by the dual-stream theory. For example, studies in cats^8^ and humans^9^ showed that spatial sensitivity in primary auditory cortex (PAC) sharpens during goal-directed sound localization, suggesting that regions that are not considered part of the spatial pathway (i.e. PAC) may nonetheless be recruited flexibly for spatial processing based on behavioral goals. Additionally, while speech processing has been attributed mostly to posterior superior temporal gyrus (STG)^10,11^, a recent study demonstrated that speech processing is instead distributed across auditory cortex^12^. While these findings indicate that sound (feature) encoding may be more distributed than posited by the hierarchical dual-stream framework, it remains unclear where the representations of these individual sound features is linked.

Additionally, it is not understood what neural mechanisms integrate cortical representations of individual sound features (e.g. spatial and non-spatial features). One hypothesis is that neuronal populations are sensitive to specific combinations of features and thereby encode multiple dimensions of an auditory object. Prior studies confirmed that some cortical sites are sensitive to multiple sound features simultaneously (e.g. in ferrets^13^, for a review^14^). However, because most prior measurements were performed with single sound sources, it is not known whether these cortical sites maintain their multi-dimensional sensitivity when presented with complex acoustic scenes comprising multiple, interfering sound sources. An alternative hypothesis states that auditory streams (pertaining to auditory objects) are formed through temporal coherence, i.e., response synchronization between neural populations that are sensitive to specific sound features^15^. Neural measurements in animals^16,17^ and humans^18^ demonstrate that temporal coherence is a plausible mechanism for auditory feature binding and segregation. It remains to be evaluated whether temporal coherence also underlies linking of voice and location features in human auditory cortex in naturalistic listening scenes.

Finally, although it is well known that auditory attention modulates the neural representation of spatial and non-spatial features^19,20^ as well as auditory object formation^21,22^, it is not known how attention modulates integrated encoding of spatial and non-spatial features in complex, naturalistic sound scenes. Moreover, it remains an open debate^2^ whether auditory objects form pre-attentively^23^ or whether attention is necessary for auditory object formation^15^.

Here, we investigated cortical multi-dimensional auditory object formation with stereotactic electroencephalography (sEEG) recordings in neurosurgical patients. We measured neural activity in response to real-world sound scenes consisting of a single localized talker or two spatially separated talkers. The unique spatiotemporal resolution of neurophysiological recordings enabled us to map feature encoding and multi-dimensional object formation across auditory cortex. We found that active listening to complex, naturalistic scenes gives rise to distributed encoding of a localized talker in auditory cortex. Cortical sites exhibit diverse tuning properties, ranging from sites that are sensitive to only location or only voice (single-feature-sensitive sites) to sites that are sensitive to both location or voice (dual-feature-sensitive sites). Furthermore, our results revealed that response patterns of dual-feature sensitive sites jointly encoded an attended talker’s voice and location features. In addition, we show that attending to a localized talker in multi-talker scenes selectively enhanced temporal coherence between single-feature sites, that is, between voice and location sensitive sites. In sum, these data demonstrate that multiple neural mechanisms contribute to linking an attended talker’s voice and location in multi-talker scenes.

## RESULTS

We analyzed neural measurements in seven neurosurgical patients recorded with intracranial depth electrodes (stereoelectroencephalography, sEEG; Methods). Three participants were implanted bilaterally, four participants were implanted in the right hemisphere only. Participants listened to English speech utterances consisting of one or two spatialized talkers. In single-talker scenes, either a male or female talker was present at a location of -45° or +45°. In two-talker scenes, a male and female talker were simultaneously present, one at -45° and the other at +45° (Figure 1 A). Speech stimuli were spatialized using a head-related transfer function (HRTF)^24^ and presented over head phones.

**Figure 1.**
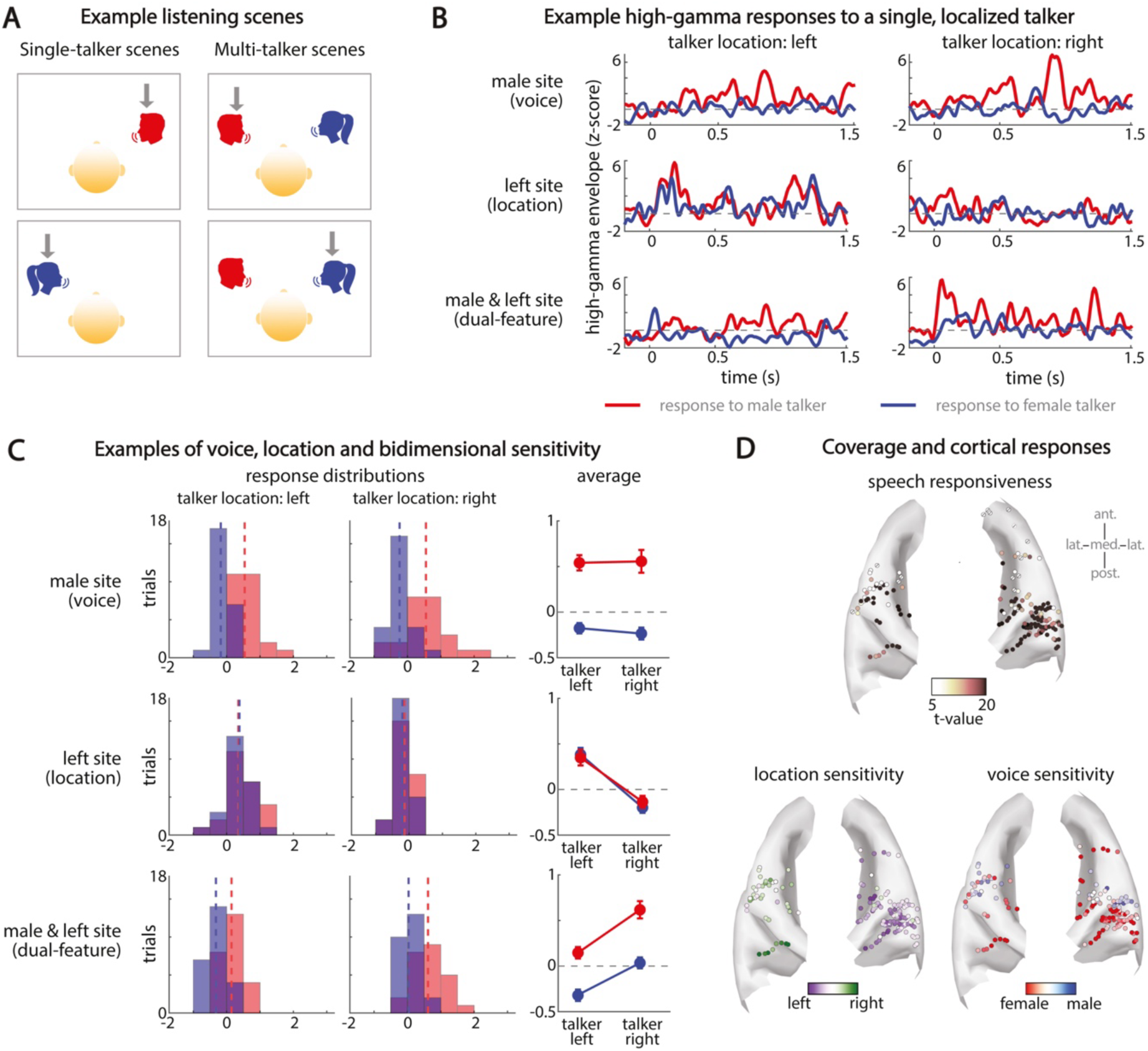
Experiment design and single-talker cortical sensitivity for location and voice features. (A) Two examples of spatial single-talker scenes (left panels) and two examples of spatial multi-talker scenes (right panels). Gray arrows indicate the attended talker. (B) Example neural responses from three sites: A single-feature voice sensitive site responding preferentially to the male talker, irrespective of the location of the talker (top panels); a single-feature location sensitive site responding preferentially to a talker on the left, irrespective of the voice of the talker (middle panels); and a dual-feature sensitive site responding preferentially to a male talker on the right (bottom panels). Sound onset was at 0 s. (C) Distribution of average trial responses to the male and female talker for the three cortical sites shown in B. Dashed line indicates the median of each distribution. Panels on the right depict the mean and standard error of the mean for each distribution. (D) Top panel: Speech responsiveness of all sites in auditory cortex. Color saturation reflects the *t*-value for the contrast speech versus silence (Methods). Sites that did not exhibit a significant response to speech are indicated by a slanted black line. Lower panels: Sensitivity for a single talker’s location (left panel) and voice features (right panel) plotted on the cortical surface for all speech responsive sites. Color indicates Cohen’s *d* (range [-1,1]).

Trials had an average duration of 5 s and the location of the talkers changed at random after each trial. The total duration of each condition (i.e., single-talker speech and multi-talker speech) was 8 minutes. For the single-talker condition, speech was paused at random intervals between trials and the participant was asked to repeat the last sentence as well as the location of the talker. For the multi-talker condition, participants were instructed at the start of a block to attend to a specific talker (i.e. ‘attend male’ or ‘attend female’). At random moments in between trials, participants were asked to report the location of the attended talker and the last sentence uttered by the attended talker. Participants successfully performed the behavioral task (mean localization accuracy = 93.8 % see^25^ for a detailed analysis of the behavioral results).

### Cortical sensitivity to a single talker’s voice and location features

We extracted the high-gamma envelope, which is thought to reflect neuronal population activity^26,27^ (Methods). We observed significant neural population responses to speech in the high-gamma envelope of 147 cortical sites in auditory cortex (paired samples t-test of responses to speech versus silence, *p* < 0.05, FDR corrected, *q* < 0.05; Figure 1 D). These speech responsive sites were located in Heschl’s gyrus (HG, 6 left hemisphere, 32 right hemisphere), planum temporale (PT, 11 left hemisphere, 24 right hemisphere) and superior temporal gyrus (STG, 25 left hemisphere, 49 right hemisphere; see Supplementary Table 1 for further details).

We characterized response properties for voice and location features by examining the responses to the single-talker scenes for each cortical site. To assess to what extent a site exhibited sensitivity to voice, to location, or to both, we contrasted the responses to one class of a feature (e.g., the male voice) to the responses to the other class of the feature (e.g., the female voice). For all sites, we extracted the mean response for each trial as the mean from 0.5 s post sound onset to 1.5 s post sound onset (that is, excluding the onset response). Figure 1 B shows example neural responses of three sites: One site sensitive to voice features (top panels, preference for male talker), one site sensitive to location features (middle panels, preference for left) and one site sensitive to both voice and location features (bottom panels, preference for male voice on the left). Figure 1 C shows the corresponding response distributions for the sites in Figure 1 B. To test for sensitivity to voice features, we computed the effect size (Cohen’s *d*^21^) for the difference between the mean responses to all male and female trials, irrespective of the location of the talker (50 trials each). To test for sensitivity to location features, we computed Cohen’s *d* for the difference between the mean responses to all trials in which the talker was at the right and all trials in which the talker was at the left, irrespective of the talker’s voice (50 trials each). Figure 1 D depicts voice and location sensitivity (Cohen’s *d*) on the cortical surface. There was no relationship between sensitivity strength for a single talker’s voice and location features (|Cohen’s *d*|, *r* = 0.037, *p* = 0.66).

Statistical testing confirmed that 47 sites were significantly sensitive to voice features only (paired samples t-tests, *p* < 0.05, FDR corrected) and 12 sites were significantly sensitive to location features only (paired samples t-tests, p < 0.05, FDR corrected). In agreement with prior results^25,28^, most sites which were sensitive to location responded preferentially to locations in the contralateral hemifield. Further, 23 sites were sensitive to both location and voice features (*p* < 0.05 for both t-tests). While multi-dimensional sensitivity has only been demonstrated for combinations of non-spatial features in humans, these results confirm prior work in animals^29^ which showed that some neuronal populations in auditory cortex are sensitive for both spatial and non-spatial features^13^. In sum, cortical responses reveal a gradient from single-feature voice or location sensitive sites to dual-feature voice and location sensitive sites.

### Spectrotemporal tuning properties explain location, voice and dual-feature sensitivity

Prior work showed that spectrotemporal tuning properties explain preferential responses to a talker’s voice^21,30^. Here, we examined whether spectrotemporal tuning properties also explain cortical location sensitivity and dual-feature sensitivity (i.e. sensitivity to both voice and location). To achieve this, we first characterized spectrotemporal tuning properties of each speech responsive site by estimating a spectrotemporal receptive field (STRF) from the site’s responses to the single-talker stimuli. We estimated STRFs using a five-fold cross-validation procedure, leaving out 20 trials and fitting the STRF on the remaining 80 trials. We used the left-out 20 trials to estimate the goodness of fit, calculating the correlation between these left-out neuronal responses and the neuronal responses predicted by the fitted STRFs (Methods). In total, 93 cortical sites had a well-fitted STRF (correlation > 0.2) and were included in the subsequent analyses.

Figure 2 A (left panel) shows the average STRF of location sensitive sites, which contains distinct response regions around 400 Hz and 600 Hz, and a more extended response region for high frequencies above ∼1 kHz. To evaluate whether the particular position of these response regions along the frequency axis of the STRF are relevant for processing location cues, we isolated spectral tuning properties by extracting the spectral receptive field (SRF) from each STRF. Specifically, the SRF corresponds to the first component of a principal component analysis of the STRF along the frequency dimension^21^ (Figure 2 A, right panel). We then calculated a correlation vector by computing for each frequency band the correlation between the SRF values of all sites at that specific frequency band and their location sensitivity (absolute Cohen’s *d*). The correlation vector (Figure 2 B, left panel) revealed high correlations for frequencies > ∼ 1.4 kHz, indicating that sites that responded strongly to high frequencies were also the most sensitive to the location of the talker.

**Figure 2.**
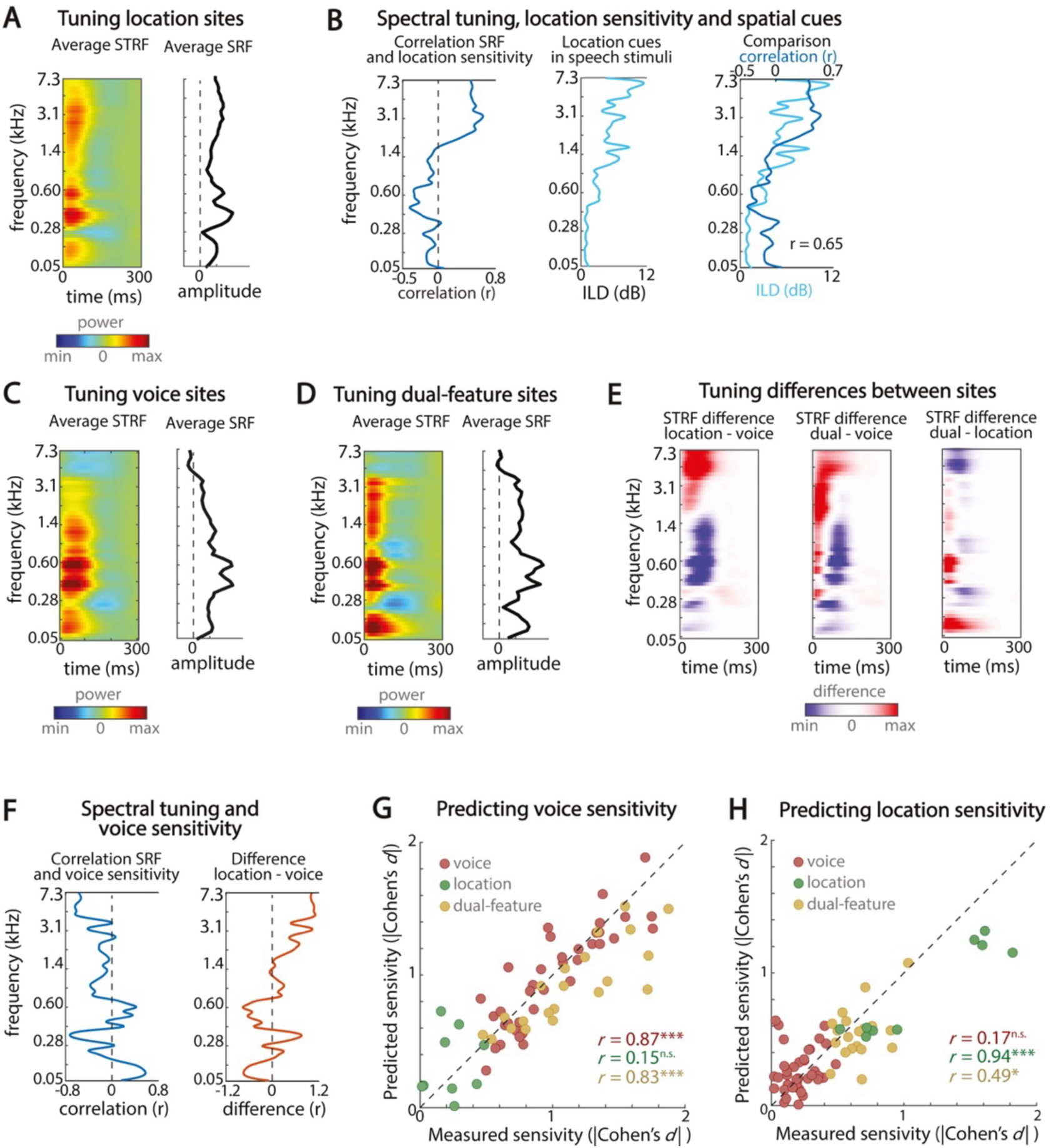
Spectrotemporal tuning characteristics explain voice, location and dual-feature sensitivity. (A) Spectrotemporal tuning properties of location sensitive sites. Left panel depicts average STRF, right panel shows the average SRF. (B) Left panel: The correlation between SRFs at each frequency band and location sensitivity. Middle panel: The presence of ILDs in the speech stimuli. Right panel: Relationship between the correlation vector and presence of ILDs. (C) Spectrotemporal tuning properties of voice sensitive sites. (D) Spectrotemporal tuning properties of dual-feature sensitive sites. (E) Comparing spectrotemporal tuning properties across voice, location and dual-feature sensitive sites. (F) Left panel: The correlation between SRFs at each frequency band and voice sensitivity. Right panel: Difference between the correlation vector for location sensitivity and for voice sensitivity. (G) Predicting voice sensitivity – that is, absolute Cohen’s *d* – from SRFs of all sites. (H) Predicting location sensitivity – absolute Cohen’s *d* – from the SRFs of all sites.

Next, we evaluated whether the spectral response profile of the location-sensitive sites can be explained by the presence of spatial cues in sounds. For humans, two sets of binaural disparity cues determine the position of a sound source in the horizontal plane: Interaural level differences (ILDs), which are most informative for frequencies > 1 kHz, and interaural time differences (ITDs), which are most informative for frequencies < 1 kHz^31^. As the temporal resolution of the high-gamma envelope – and therefore also the resolution of the STRFs – is too low to capture responses at the ITD timescale (that is, < 1 millisecond), we focus on ILDs in our analysis. We calculated ILD for each stimulus as the absolute difference in power between the spectrum of the left and right channel, expressed in decibel (dB). Figure 2 B (middle panel) shows that ILDs were most pronounced in higher frequencies, especially above ∼1.4 kHz. Crucially, Figure 2 B (right panel) shows that there was a strong relationship between the correlation vector and the presence of spatial cues in the speech stimuli (*r* = 0.65, *p* ≪ 0.0001). Thus, as a consequence of the high-frequency tuning of location-sensitive sites, ILDs within these high frequency components strongly modulate the responses of these sites, leading to a robust cortical representation of sound location.

We then assessed to what extent the spectrotemporal tuning properties of location-sensitive sites differ from the tuning properties of voice sensitive sites and from dual-feature-sensitive sites. Figure 2 C and D show the average STRF and SRF for voice and dual-feature-sensitive sites, respectively. The STRFs of voice-sensitive sites exhibited strong responses especially at frequencies below 1.4 kHz. Moreover, in line with previous work^21,25,30^, the STRFs of voice-sensitive sites exhibited spectrotemporal tuning characteristics matching the acoustic profile of the preferred talker (Supplementary Figure 1). The STRFs of dual-feature-sensitive sites exhibited similar response regions < 1.4 kHz, but contained an additional extended response region between 1 kHz and 3 kHz. Figure 2 E visualizes the difference between the average STRFs of the location-, voice- and dual-feature-sensitive sites. These plots show that both the location- and dual-feature-sensitive sites exhibit stronger responses to frequencies > 1 kHz than voice-sensitive sites (left and middle panel). Furthermore, while responses of dual-feature-sensitive sites to frequencies between ∼1 kHz and ∼3 kHz were comparable to responses of location-sensitive sites, their tuning properties differed in low frequencies (< ∼700 Hz) and in the highest frequencies. Specifically, dual-feature-sensitive sites showed stronger responses to low frequencies than location-sensitive sites, but weaker responses to frequencies > ∼ 3.5 kHz.

Next, similar to our analysis of location sensitivity across cortical sites (Fig. 2 B), we evaluated to what extent spectral tuning properties are related to cortical voice sensitivity by calculating for each frequency band the correlation between SRF and voice sensitivity (absolute Cohen’s *d*). Figure 2 F (left panel) depicts the resulting correlation vector, which shows that sites with strong responses to frequencies < ∼200 Hz and to frequencies between ∼400 Hz and ∼700 Hz were more sensitive to voice (positive correlation), while sites with strong responses to frequencies > ∼3.5 kHz were less sensitive to voice (negative correlation). Fig. 2 F (right panel) shows the difference between the correlation vector for location sensitivity (Fig. 2 B) to the correlation vector for voice sensitivity. This demonstrates that responses to high frequencies are mostly related to location sensitivity, while responses to distinct low frequency bands (< ∼1 kHz) are mostly related to voice sensitivity.

Finally, to quantify the relationship between a site’s spectral tuning properties and sensitivity to a talker’s location and a talker’s voice, we mapped SRFs to location and to voice sensitivity using ridge regression (Methods). Fig. 2 G shows that when SRFs are mapped to voice sensitivity, the regression model can accurately predict voice sensitivity of voice-sensitive sites (*r* = 0.87, *p* ≪ 0.0001) as well as of dual-feature-sensitive sites (*r* = 0.83, *p* ≪ 0.0001), but not of location-sensitive sites (*r* = 0.15, *p* > 0.05). In contrast, when SRFs are mapped to location sensitivity, the regression model accurately predicts location sensitivity of location-sensitive sites (*r* = 0.94, *p* = 0.0001) as well as dual-feature-sensitive sites (*r* = 0.49, *p* = 0.025), but not of voice-sensitive sites (*r* = 0.17, *p* > 0.05; Fig. 2 H). These findings demonstrate that the regression model learns a different set of weights for mapping SRF to voice sensitivity than for mapping SRF to location sensitivity, highlighting that voice and location sensitivity can be attributed to spectral tuning properties (see Supplementary Figure 2 for the learned sets of weights). Further, these results confirm that dual-feature sensitive sites share spectral tuning properties with both location- and voice-sensitive sites.

Taken together, we demonstrate that location-, voice- and dual-feature-sensitive sites exhibit distinct spectrotemporal tuning properties and that these tuning properties explain their sensitivity to a talker’s voice, location, or a combination thereof.

### Sensitivity to a talker’s voice and location across the cortical hierarchy

To investigate to what extent sensitivity to a talker’s voice and location can be related to cortical processing stages, we investigated how sensitivity to a talker’s features was distributed across auditory cortex. While several studies linked delineated anatomical regions to hierarchical processing stages (for example, HG is considered primary auditory cortex and PT and STG higher-order auditory regions^30^), other work investigating neural response latencies and response properties showed that a single anatomical region may contain different auditory processing stages^12,32^. That is, as response latency roughly corresponds to the number of synapses away from the periphery, response latency is considered an indication of the processing stage of a neural site. Here, we assessed the distribution of feature sensitivity both within cortical auditory regions and as a function of response latency. We calculated response latency as the peak along the temporal dimension of the STRF (for sites with a well-fitted STRF, *r* > 0.2, *n* = 93; Methods).

Figure 3 A shows the regional distributions of voice sensitivity (Cohen’s *d*). Comparing the distributions across regions showed that sensitivity to a talker’s voice was stronger in HG than in STG (|Cohen’s *d*|, Kruskal-Wallis H test, χ^2^(2) = 14.6, *p* = 0.0007, Figure 3 A). Further, there was a negative correlation between sensitivity to a talker’s voice and response latency (*r* = -0.53, *p* = 1.3E-7; Figure 3 B, left panel). These findings confirm prior reports of a decrease in sensitivity to a talker’s voice along the cortical auditory processing hierarchy^21^. In contrast, although we observed a trend towards regional differences in location sensitivity (|Cohen’s *d*|, Kruskal-Wallis H test, χ^2^(2) = 5.45, *p* = 0.07; Figure 3 A), this trend failed to reach significance. Moreover, we observed no correlation between sensitivity to a talker’s location and response latency (*r* = -0.15, *p* = 0.16; Figure 3 B, right panel). While the lack of regional differences may be attributed to the relatively low anatomical sampling density, the response latency results also indicate that location sensitivity is consistent across low- and high-level processing stages during active listening. These findings confirm recent work^9,25^, but contrast the predictions of the dual-stream framework which posits that PT is functionally specialized for spatial processing^6,7,33^.

**Figure 3.**
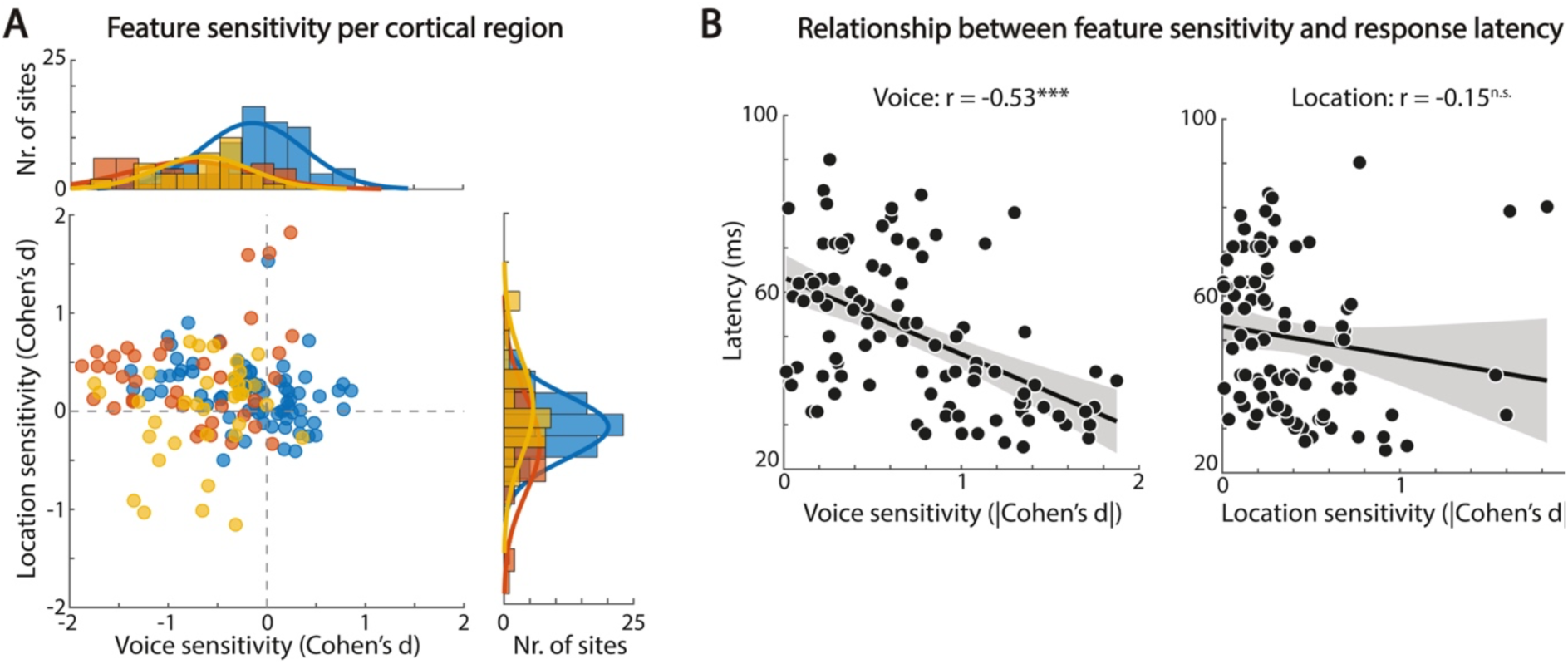
Sensitivity to a single talker’s voice and location across the cortical hierarchy. (A) Scatterplot of voice sensitivity (x-axis) and location sensitivity (y-axis). Each symbol represents an individual site. Bar graphs depict corresponding marginal distributions for voice sensitivity (left) and location sensitivity (right). (B) Correlation between single-talker response latency and feature sensitivity (left panel: voice; right panel: location). Each circle depicts a site. Solid lines depict the correlation; shaded areas depict the 95% confidence interval. Asterisks indicate significance.

### Attentional modulation of response gain by a talker’s voice and location in multi-talker scenes

We showed that cortical sites exhibit varying degrees of sensitivity for a single talker’s location and voice. Motivated by prior reports about the attentional modulation of neuronal responses to a talker’s voice and location^21,25^, we examined to what degree we observed such attentional response gain modulations in multi-talker scenes in our data. Further, we characterized how attentional response gain modulation by a talker’s voice relates to attentional response gain modulation by a talker’s location. The analysis of attentional response gain modulation in multi-talker scenes was similar to the quantification of sensitivity to a single talker’s voice and location features: We calculated the effect size (Cohen’s *d*) for the difference between the mean response across trials for each attentional condition. For example, we calculated Cohen’s *d* for the difference between the mean response across trials in which the male talker was attended and the mean response across trials in which the female talker was attended. As before, we calculated the mean response for each trial from 0.5 s post sound onset to 1.5 s post sound onset, excluding the onset response.

In agreement with prior studies^21,25^, attending a localized talker evoked weak response gain modulations across speech responsive sites, both by the attended talker’s voice and by the attended talker’s location (Figure 4 A). Statistical testing did not identify neural sites which exhibited significant attentional response gain modulation either by the attended talker’s voice (paired samples t-tests, *p* > 0.05), location (paired samples t-tests, *p* > 0.05), or both combined (*p* >0.05). Further, Figure 4 B shows that attentional response gain modulation by the attended talker’s voice or location was significantly weaker than single-talker voice or location sensitivity (paired samples t-test of |Cohen’s *d*|; voice: *t*(146) = 9.65, *p* = 2.38E-17; location: *t*(146) = 6.68, *p* = 4.64E-10). Single-talker voice sensitivity was not correlated to multi-talker attentional response gain modulation by the attended talker’s voice (*r* = 0.13, *p* = 0.11), while single-talker location sensitivity was moderately correlated to multi-talker attentional gain modulation by the attended talker’s location (*r* = 0.40, *p* ≪ 0.0001).

**Figure 4.**
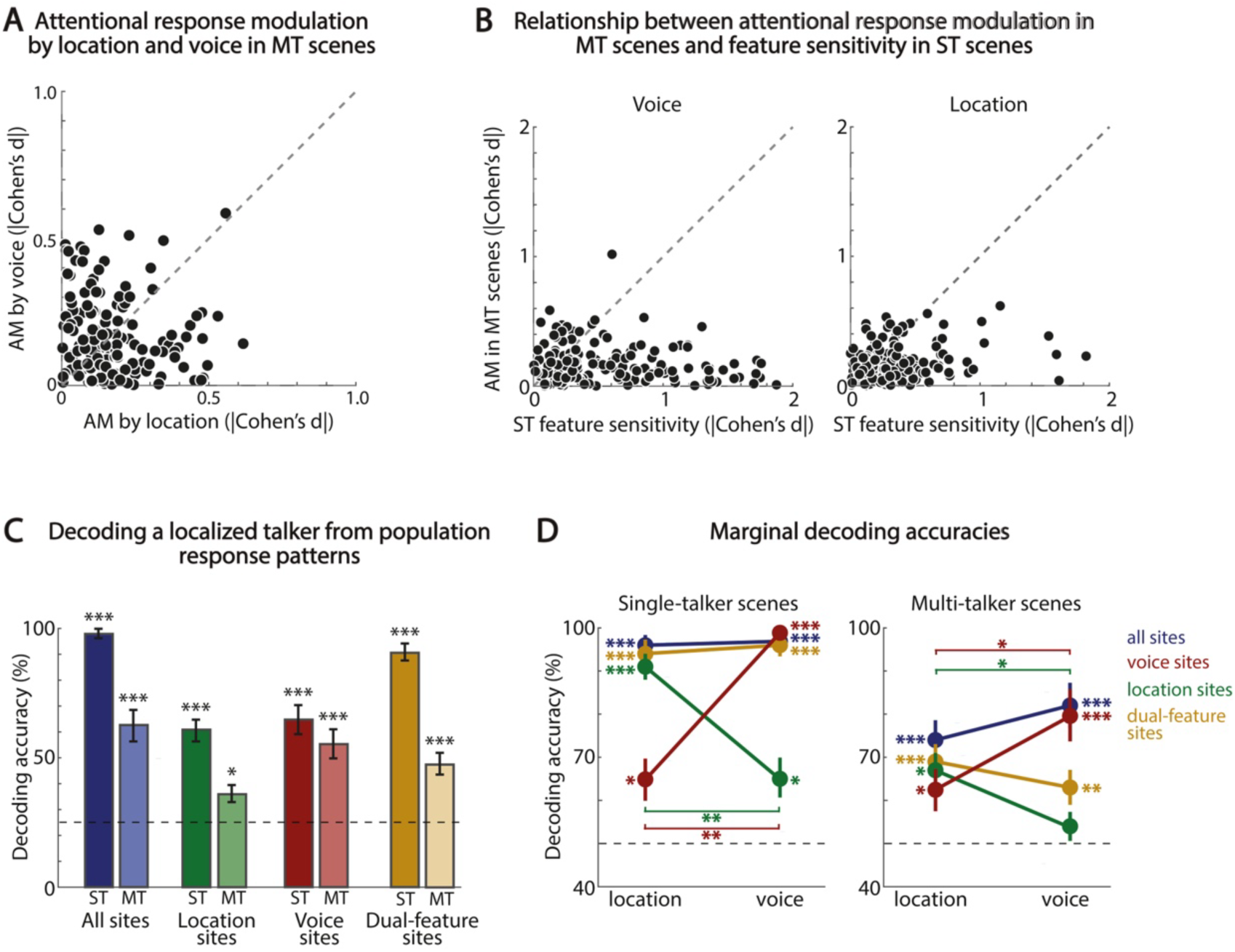
Attentional response gain modulation and the representation of a localized talker in population response patterns in single- and multi-talker scenes. (A) Scatterplot of response gain modulation by attending to a talker’s location (x-axis, |Cohen’s *d*|) and by attending to a talker’s voice (y-axis, |Cohen’s *d*|). (B) Relationship between attentional response gain modulation in multi-talker scenes and single-talker sensitivity. Left panel: Single-talker voice sensitivity (x-axis) versus multi-talker response gain modulation by attending to a talker’s voice (y-axis). Right panel: Single-talker location sensitivity (x-axis) versus multi-talker response gain modulation by attending to a talker’s location (y-axis). (C) Decoding a localized talker from response patterns in single-talker scenes (ST) and decoding an attended localized talker in multi-talker scenes (MT). Horizontal lines depict chance level. Asterisks indicate significance: *** = *p* < 0.001. (D) Marginal decoding accuracies for a talker’s voice and location. Dashed line depicts chance level, asterisks indicate significance: * = p < 0.05; ** = p < 0.01; *** = p < 0.001.

These findings are in agreement with prior work showing that sites which exhibit a preference for a particular voice in single-talker scenes do not exhibit similar attentional response gain modulation in multi-talker scenes^21^ and raise questions about the precise role of ‘functionally specialized’ sites in multi-talker scenes. That is, neuronal sensitivity to a single sound source’s features (e.g. pitch, location) is generally considered an indication of functionally specialized processing^2,33^. However, as these sites do not exhibit corresponding, dedicated attentional response gain modulation based on the attended sound source’s features in multi-source scenes, it appears that the functionally specialized responses observed in single-source scenes are not sustained in naturalistic multi-source scenes.

### Decoding a localized talker from population activity patterns in multi-talker scenes

To elucidate the relationship between the encoding properties of individual cortical sites and population encoding properties, we examined whether a localized talker can be decoded from population response patterns. Specifically, we used a linear decoding approach to assess to what extent a localized talker can be decoded from population responses in single talker scenes and to what extent an attended localized talker can be decoded from population responses in multi-talker scenes. To decode a localized talker in single-talker scenes, we trained a four-class regularized least-squares (RLS^22,34^) classifier on the response patterns in single-talker scenes using a leave-two-trials-out cross-validation procedure (corresponding to 25 folds). To decode an attended localized talker, we trained an identical four-class RLS classifier on the response patterns in multi-talker scenes using a similar cross-validation procedure. We assessed decoding accuracy by predicting the (attended) talker’s voice and location from the response patterns of the left-out trials of each fold (Methods).

As expected, Figure 4 C shows that a single localized talker could be accurately decoded from the entire population of speech responsive sites (*n* = 147; average accuracy [standard error of the mean; SEM] = 93.0 % [2.29], *p* = 0, FDR corrected). Similarly, the attended localized talker was decoded accurately from the entire population of speech responsive sites (mean accuracy [SEM] = 63.0 % [5.80], *p* = 0). The marginal decoding accuracies for the talker’s voice and location in Figure 4 D show that both features were decoded with equal accuracy in single-talker scenes (mean marginal accuracy: voice [SEM] = 97.0 % [1.66], *p* = 0; location [SEM] = 96.0 % [1.87], *p* = 0; paired samples *t*-test, *t*(24) = 0.37, *p* = 0.72) as well as in multi-talker scenes (mean accuracy voice [SEM] = 82.0 % [4.68], *p* = 0; average accuracy location [SEM] = 74.0 % [3.95], *p* = 0; *t*(24) = 1.69, *p* = 0.14). These findings show that despite the weak attentional modulation of responses of individual cortical sites by the attended talker’s voice and location in multi-talker scenes (Figure 4 A, B), response patterns across the entire population of speech responsive sites nevertheless represented the attended localized talker with high fidelity.

Next, we examined how sites which exhibit single-feature sensitivity for a talker’s voice or location features in their local responses (*n* = 47 and *n* = 12, Figure 1) encode a localized talker in their population response patterns. We therefore trained the RLS classifier on the population responses of these neural sites with the same procedure described above. Note that although the latter population is relatively small, we chose to use this stringent selection to ensure that the population did not incorporate sites that were also to some extent sensitive to a talker’s voice. Figure 4 C shows that the classifier successfully decoded a localized talker in single-talker scenes from voice sensitive sites (mean accuracy [SEM] = 65.0 % [4.33], *p* = 0) as well as from location sensitive sites (mean accuracy 61.0 % [4.10], *p* = 0). However, the marginal accuracies in Figure 4 D show that the classifier decoded the talker’s voice and location features with different accuracy from these populations. Specifically, the talker’s voice was decoded more accurately from population responses of voice sensitive sites than the talker’s location (mean marginal accuracy: voice [SEM] = 99.0 % [1.00], *p* = 0; location [SEM] = 65.0 % [4.33], *p* = 0.0098; paired samples t-test, *t*(24) = 7.49, *p* = 3.93E-7). Conversely, decoding accuracy was higher for the talker’s location than for the talker’s voice when the classifier operated on population responses of location sensitive sites (voice [SEM] = 65.0 % [4.33], *p* = 0.02; location [SEM] = 91.0 % [2.45], *p* = 0.0024; *t*(24) = 5.32, *p* = 3.74E-5). These findings show that the population responses of single-feature sensitive sites encode their preferred feature dimension with high fidelity but also encode coarse information about other feature dimensions of an individual talker.

In multi-talker scenes, the classifier decoded the attended localized talker above chance level from population responses of voice sensitive sites (Figure 4 C, mean accuracy [SEM] = 55.0 % [5.59], *p* = 0) as well as from population responses of location sensitive sites (Figure 4 C, mean accuracy [SEM] = 36.0 % [3.84], *p* = 0.026). However, while both the attended talker’s location and voice were decoded above chance level from population responses of voice sensitive sites (mean marginal accuracy: voice [SEM] = 80.0 % [5.40], *p* = 0; location [SEM] = 63.0 % [4.59], *p* = 0.023), only the attended talker’s location was decoded accurately from population responses of location sensitive sites. Specifically, the decoding accuracy for the attended talker’s voice just failed to reach statistical significance, which may be a consequence of the small number of sites in this group (mean marginal accuracy: voice [SEM] = 54.0 % [2.77], *p* = 0.079; location [SEM] = 67.0 % [3.45], *p* = 0.021). For both voice and location sensitive sites, the preferred feature was decoded significantly better than the other feature (voice: *t*(24) = 2.72, *p* = 0.024; location: *t*(24) = 2.98, *p* = 0.024). These findings indicate that populations consisting of cortical sites which exhibit tuning properties that resemble functional specialization in response to single-source sound scenes (that is, single-feature sensitive sites) may nonetheless encode (coarse) information about other dimensions of an attended auditory object in multi-talker scenes. Future work including more fine-grained sampling of multiple feature dimensions (e.g., more talkers and more voices) is required to establish the resolution with which population responses of single-feature sensitive sites encode other feature dimensions in multi-talker scenes.

Finally, we showed previously that some cortical sites were sensitive both for a talker’s voice and location, that is, dual-feature sensitive sites (Figure 1, *n* = 23). We examined to what extent the population response patterns of these sites jointly encode a talker’s voice and location in single- and in multi-talker conditions. Figure 4 C shows that the localized talker was decoded accurately from population responses of dual-feature sensitive sites in single-talker scenes (mean accuracy [SEM] = 91.0 % [3.50], *p* = 0) as well as from population responses in multi-talker scenes (mean accuracy [SEM] = 47.0 % [3.63], *p* = 0). Importantly, Figure 4 D shows that dual-feature sensitive encoded the talker’s voice and location features with equal accuracy, both in single-talker (mean marginal accuracy: voice [SEM] = 96.0 % [1.87], *p* = 0; location [SEM] = 94.0 % [2.61], *p* = 0; paired samples t-test: *t*(24) = 0.81, *p* = 0.57) and in multi-talker scenes (mean marginal accuracy: voice [SEM] = 63.0 % [3.57], *p* = 0.006; location [SEM] = 69.0 % [3.62], *p* = 0; paired samples t-test: *t*(24) = 1.1, *p* = 0.28). In sum, we show that population responses of dual-feature sensitive sites encode both spatial and non-spatial features of an attended talker in multi-talker scenes. This suggests that such dual-feature sensitive sites contribute to the encoding of multiple dimensions of an auditory object.

### Sites which selectively track a single, attended speech stream in multi-talker scenes simultaneously encode the attended talker’s voice and location features

In the preceding sections, we examined to what extent the population response patterns sites that exhibit sensitivity to a single feature (voice, location) or both features (voice and location, dual-feature sensitive) in their responses to a single talker, represent an attended talker’s voice and location features in multi-talker scenes. However, prior work showed that auditory cortex also contains neural sites which are not strongly sensitive to a single talker’s features, but which nonetheless play a crucial role in auditory object formation by selectively tracking a single, attended speech stream in multi-talker listening scenes^21,22,24,34^. As the relationship between such speech stream tracking and encoding of the attended talker’s features is not known, we analyzed the neuronal responses in multi-talker scenes to evaluate to what degree cortical sites which selectively track a single, attended speech stream additionally encode the voice and location features of the attended talker. Furthermore, while it has been demonstrated that attentional modulation of spectrotemporal tuning properties^35^ forms the physiological basis for such selective tracking of speech^25^, the link between selective tracking of the attended speech stream, attentional modulation of spectrotemporal tuning properties, and encoding of an attended talker’s location and voice features is presently unclear.

For each site, we first quantified selective tracking of a single, attended speech stream by calculating to what extent a site’s responses to spatial multi-talker scenes were modulated by attention such that response time courses in multi-talker scenes resembled response time course to that same localized talker in single-talker scenes. That is, we define the tracking index (TI) for each site similar to the definition in ^21^:

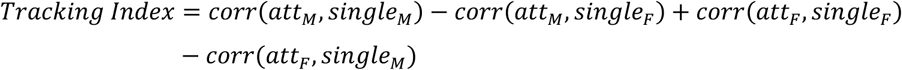

Here, *M* refers to the male talker and *F* to the female talker. Further, *corr*(*att, single*) corresponds to the correlation between the response time course when attending an individual talker in multi-talker scenes and the response time course to that individual talker in single-talker scenes. That is, as multi-talker scenes were constructed by combining two single-talker scenes, we had measures of responses to the same speech stream in single-talker and in multi-talker conditions (see example in Figure 5 A). A high TI value indicates that the response time course in the multi-talker scene resembles the response time course to that talker in isolation in the single-talker scenes. Figure 5 B shows that TI varied across cortical sites, with approximately ∼33 of the sites exhibiting a TI ≥ 0.1. Note that sites which exhibited selective tracking of a single, attended talker in multi-talker scenes (i.e., sites with a high TI) were distinct from feature sensitive sites (correlation between TI and single-talker voice sensitivity: *r* = 0.11, *p* = 0.24; correlation between TI and single-talker location sensitivity: *r* = 0.12, *p* = 0.24). Furthermore, TI was not related to attentional response gain modulation by the attended talker’s features either (correlation TI and attended talker’s voice: *r* = 0.06, *p* = 0.48; correlation TI and attended talker’s location: *r* = -0.13, *p* =0.24).

**Figure 5.**
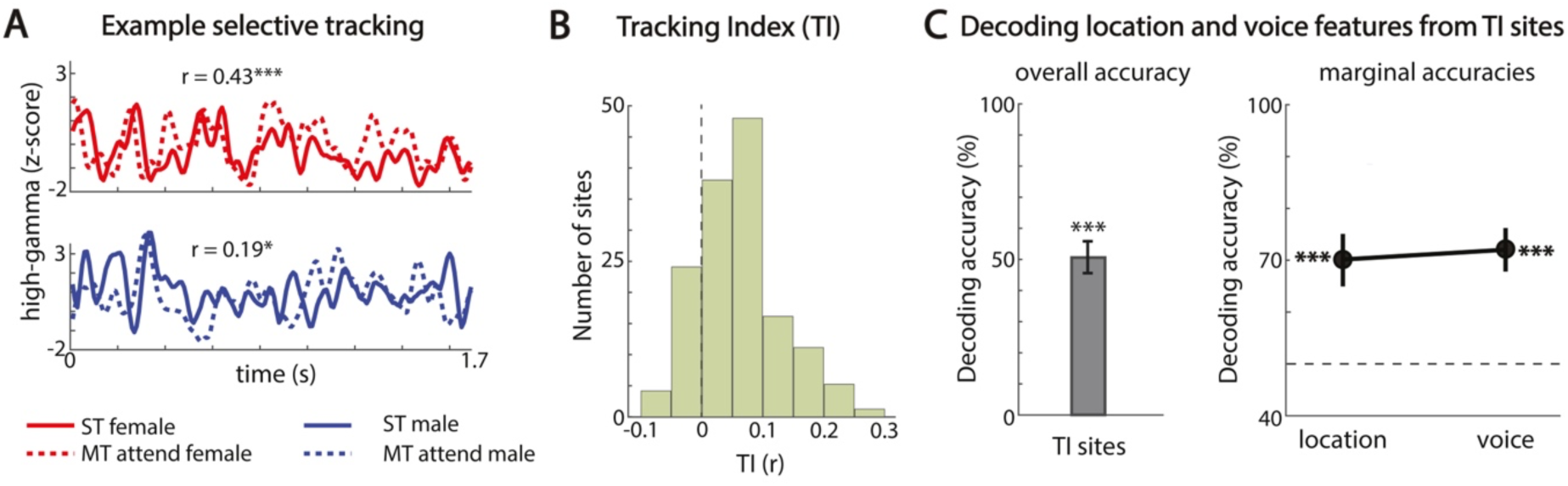
Selective speech tracking and encoding of the attended talker’s voice and location. (A) High-gamma responses of an example site exhibiting selective tracking of the attended talker. Solid lines depict response in single-talker scene, dotted lines depict response in multi-talker scene. (B) Distribution of selective tracking indices across cortical sites. (C) Left panel: Decoding an attended localized talker from population responses in multi-talker scenes. Horizontal line depicts chance level. Asterisks indicate significance: *** = *p* < 0.001. Right panel: Marginal decoding accuracies for a talker’s voice and location. Dashed line depicts chance level, asterisks indicate significance: * = p < 0.05; ** = p < 0.01; *** = p < 0.001.

Next, we examined to what extent the population response patterns of sites with a high TI – that is, sites which selectively track a single, attended speech stream – also encode the attended talker’s voice and location features. We trained the four-class classifier on population response patterns in multi-talker scenes of sites with a high TI (i.e., for sites with TI > 0.1, *n* = 33). The classifier accurately decoded the attended localized talker from these population response patterns (average accuracy [SEM] = 51.0 % [5.10], *p* = 0; Figure 5 C). Furthermore, the classifier decoded the attended talker’s voice and location with equal accuracy (Figure 5 D; marginal accuracies: voice [SEM] = 72.0 % [3.63], *p* = 0; location [SEM] = 70.0 % [4.56], *p* = 0; paired samples t-test, *t*(24) = 0.40, *p* = 0.69). In sum, population response patterns of sites which selectively tracked a single, attended speech stream also encoded the attended talker’s voice and location features. These findings thus indicate that population responses of such sites play a crucial role in combining selective tracking of an attended auditory object (here, speech stream) with encoding of the features of that object (here, the talker’s voice and location).

### Attention to a localized talker in multi-talker scenes enhances temporal coherence between voice- and location-sensitive sites

We showed that an attended talker’s voice and location features are linked by population response patterns which simultaneously encode an attended talker’s voice and location features in spatial multi-talker scenes. However, other mechanisms may also contribute to linking an attended talker’s voice and location features. In particular, it has been proposed that attending to a particular sound object enhances binding of its features by increasing the temporal coherence of the slow fluctuations in the neuronal responses of sites that are sensitive to a single sound feature^15^. We therefore evaluated to what extent temporal coherence contributed to linking the attended talker’s voice and location features in the present experiment.

We computed temporal coherence between neural responses in multi-talker scenes for pairs of voice- and location-sensitive (i.e., based on their single-talker sensitivity to voice or location, Fig. 1). For each voice-location pair (*n* = 57), we quantified temporal coherence of the high gamma envelope at frequencies between 2-22 Hz using the coherency coefficient. The coherency coefficient is the frequency-domain mathematical equivalent of the cross-correlation function in the time-domain^36^ (within-subjects analysis; Methods). We evaluated the hypothesis that attention selectively enhances temporal coherence between the voice- and location-sensitive site in each pair by contrasting temporal coherence across attention conditions. That is, we examined whether temporal coherence increased when attention was directed towards a localized talker which matched the pair’s preferred features (condition ‘preferred features attended’) in comparison to when attention was directed to a localized talker which was orthogonal to the pair’s preferred features (condition ‘preferred features unattended’):

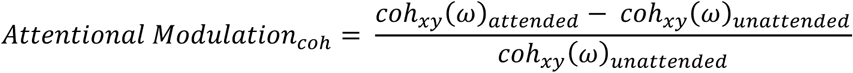

Here, *xy* refers to a pair of sites consisting of one location-sensitive site (*x*) and one voice-sensitive site (*y*), while ω refers to the frequency range under consideration. For example, for a pair of sites consisting of a voice-sensitive site tuned to the *female talker* and a location-sensitive site tuned to the *right*, we hypothesized that temporal coherence increases when attention is directed to a female talker on the right in comparison to when attention is directed to a male talker on the left (Figure 6 A).

**Figure 6:**
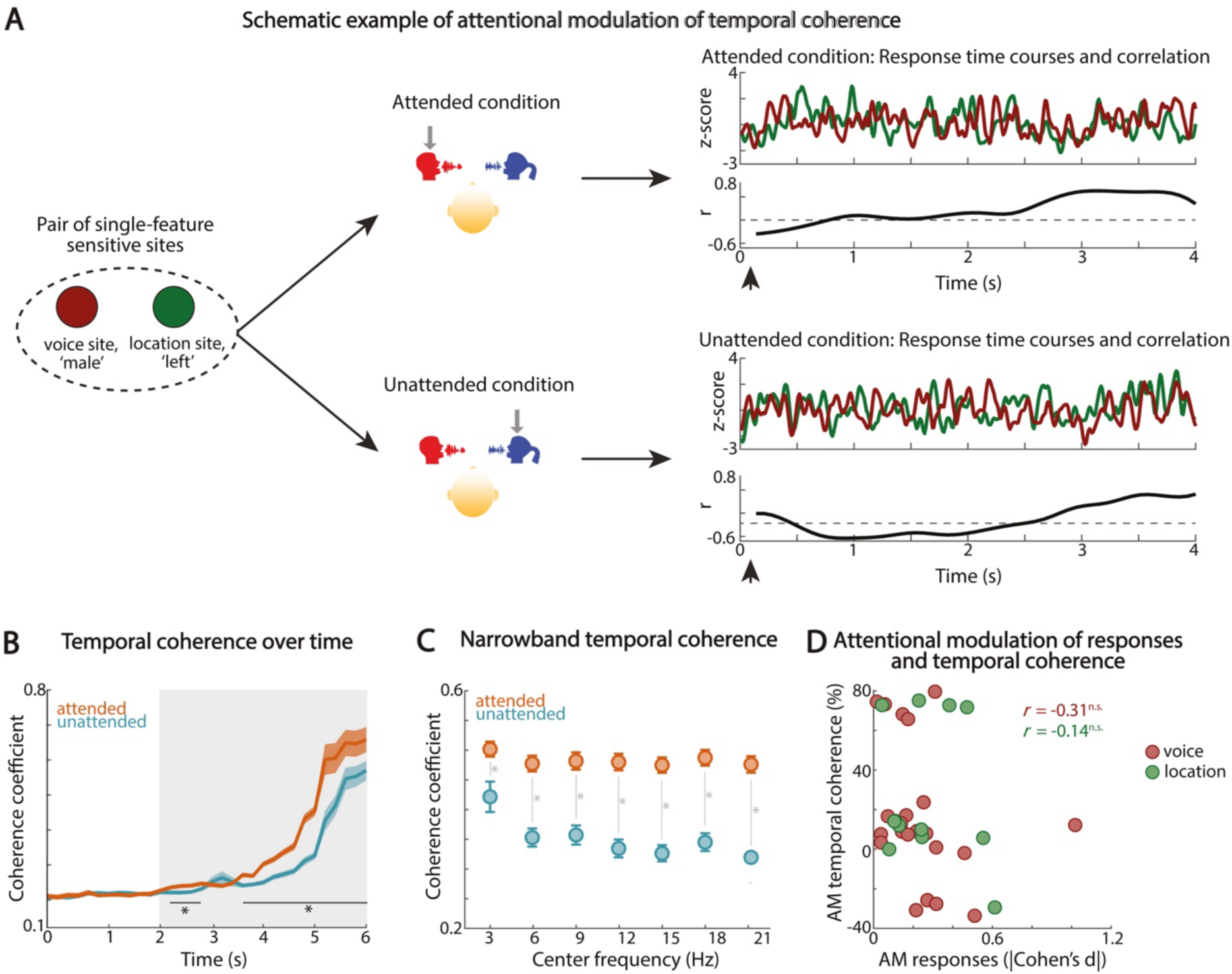
Linking an attended talker’s voice and location through temporal coherence. (A) Example of attentional modulation of the correlation between the high-gamma response envelopes of a pair of neural sites consisting of a voice sensitive site (brown) and a location sensitive site (green). When attention is directed to the preferred features of the pair of sites, the correlation between the high-gamma envelope increases over time (right panel, top). When attention is directed to the non-preferred features, the increase in correlation is much lower (right panel, bottom). (B) Development of average broadband temporal coherence (2-22 Hz) over time when attention is directed towards their preferred features (red) and when their preferred features are unattended (blue). Shaded area reflects standard error of the mean (SEM). Horizontal lines indicate time points at which the coherence coefficient is significantly different between attended and unattended conditions. The gray shaded area indicates the time-scale of perceptual reports of stream formation^37,38^. (C) Temporal coherence per narrowband frequency bin. Vertical lines indicate at which frequency bins the coherence coefficient is significantly different between the attended and unattended condition. (D) Bars show the average strength for single-talker feature sensitivity (dark bars) and multi-talker attentional gain modulation (gray bars). Asterisks indicate a significant difference between single-talker sensitivity and multi-talker attentional gain modulation. (D) Scatterplot depicts attentional modulation (AM) of responses (x-axis) and temporal coherence (y-axis) for voice sensitive sites (red circles) and location sensitive sites (green circles).

To begin with, we evaluated the development of attentional modulation of temporal coherence over time by calculating broadband temporal coherence (i.e. across all frequencies between 2 and 22 Hz) in shifting windows of 1,000 ms with a 200 ms hop length. Figure 6 B shows that the temporal coherence between voice- and location-sensitive sites starts to rise after 2 seconds, both when attention is directed towards their preferred features (that is, attended condition) and when attention is directed towards their non-preferred features (that is, unattended condition). However, the temporal coherence between voice-location-sensitive sites increases more when attention is directed towards the preferred features: The coherence coefficient is significantly higher in the attended than in the unattended condition from 2.2 s onwards (one-sided permutation test, *p* < 0.05, FDR corrected; Figure 6 B). This time scale is in line with the perceptual experience of auditory stream formation, as listeners typically start to perceive two segregated streams a few seconds after the onset of two simultaneous sequences (e.g. ^37,38^).

Next, we examined to what extent the observed attentional enhancement of temporal coherence over time was frequency specific. We therefore repeated the analysis, but using only the late response which showed the most robust attentional temporal coherence gain (i.e., from 2 until 6 s post sound onset) and evaluating coherence for narrow frequency bins of 3 Hz (central frequencies [CF]: 3, 6, 9, 12, 15, 18, 21 Hz). Figure 6 C shows that temporal coherence between voice- and location-sensitive sites was higher in the attended than in the unattended condition in all frequency bins (one-sided permutation test, *p* < 0.05, FDR corrected). Further, there was a significant difference in temporal coherence between frequency bins in the unattended condition (repeated measures ANOVA, *F*(6,336) = 10,18, *p* = 2.34E-10; Figure 6 C). Post-hoc multiple comparisons demonstrate that temporal coherence is higher for frequency bin with CF = 3 Hz than for any other frequency bin (*p* < 0.05, Tukey-Kramer correction for multiple comparisons). As this frequency bin corresponds to the syllabic rate, these findings suggest that the observed temporal coherence between voice and location sites in the 2 – 4 Hz frequency range may be driven by neural synchronization to the syllabic rate^39–42^. In contrast, we did not observe a difference in temporal coherence between frequency bins in the attended condition (repeated measures ANOVA, *F*(6,336) = 0.84, *p* = 0.54). Taken together, these results indicate that attention enhances temporal coherence between the high-gamma envelopes of voice- and location-sensitive sites across all frequencies between 2-22 Hz, and that this attentional modulation is largest for frequencies ≥ 5 Hz.

Finally, we evaluated the relationship between the attentional modulation of temporal coherence and attentional response gain modulation to ensure that attentional enhancement of temporal coherence is not solely attributable to the attended talker dominating neural responses. Figure 6 D shows that there was no correlation between attentional modulation of temporal coherence and attentional response gain modulation either for voice- or location-sensitive sites, indicating that attentional response gain modulation is unlikely to drive attentional modulation of temporal coherence. Additionally, we examined whether attentional modulation of temporal coherence between pairs of location- and voice-sensitive sites was a consequence of selective tracking of a single, attended speech streams (i.e., see Figure 5 for selective tracking). Yet, our data showed that location- and voice-sensitive sites had a low tracking index (TI): average TI [SEM] for location-sensitive sites was 0.057 [0.005] and for voice-sensitive sites 0.064 [0.003]. Taken together, these findings indicate that the attentional enhancement of temporal coherence between voice- and location-sensitive sites was independent of other attentional effects on neural responses, such as attentional response gain modulation and selective tracking of a single, attended speech stream.

In sum, our results demonstrate that temporal coherence between voice- and location-sensitive sites increases over time in spatial multi-talker scenes, both with and without attention to the preferred features, but the increase in temporal coherence is significantly stronger in the attended condition. These findings indicate that temporal coherence is a plausible binding mechanism for linking voice and location encoding by single-feature sensitive sites in order to form a complete auditory object in complex, multi-source auditory scenes.

## DISCUSSION

In real-world environments, listeners must navigate complex acoustic scenes to extract meaningful information from overlapping sound sources, a process that relies on the formation of multi-dimensional auditory objects. This study investigated how the human auditory cortex integrates distinct features—voice and spatial location—of an attended talker during multi-talker scenes. Using invasive intracranial recordings, we demonstrated that auditory object formation involves two complementary neural mechanisms: joint population coding and temporal coherence. First, we found that cortical responses exhibit a gradient of feature sensitivity, ranging from single-feature-sensitive sites (responsive primarily to voice or location) to dual-feature-sensitive sites (responsive to both). At the population level, response patterns from both single- and dual-feature-sensitive sites jointly encoded the attended talker’s voice and location, highlighting the distributed and multi-dimensional nature of feature representation in the auditory cortex. Importantly, even single-feature sensitive sites encoded coarse information about secondary features, underscoring the flexibility of population coding mechanisms. Second, we showed that attention selectively enhanced temporal coherence between voice- and location-sensitive sites. This enhancement was independent of attention-driven response gain, indicating that temporal coherence reflects a distinct mechanism for feature integration. Together, these findings suggest that multi-dimensional auditory object formation emerges from a combination of joint coding and attention-driven synchronization, enabling robust representation of an attended talker in complex, naturalistic listening environments.

### Active task design and naturalistic stimuli reveal distributed voice and location encoding

Our data showed that the sensitivity of local cortical sites for a talker’s voice and location features can be explained by the underlying spectrotemporal tuning properties. These results align with prior research attributing speaker sensitivity to spectrotemporal tuning properties^21^ and fast temporal processing to the posterior-dorsal regions of human auditory cortex^43^ which tend to show strong spatial sensitivity^28^. Additionally, our findings indicate that cortical representations of a multi-dimensional localized talker are derived from joint encoding in distributed population response patterns rather than separate voice and location encoding in dual processing streams within delineated anatomical regions^6,7,33^. Moreover, linking local responses to population encoding showed that sites which are characterized by functionally specialized local responses (for example, voice sensitive sites), nevertheless encode information about both voice and location in their population responses.

Further, the distributed networks of voice and location sensitivity that we observed at the level of individual cortical sites is in agreement with a recent study which demonstrated that acoustic and phonetic processing in auditory cortex are based on distributed, parallel processing rather than serial processing^12^. This indicates that distributed processing may be a general characteristic of auditory encoding and speech encoding specifically^10^. Moreover, the occurrence of distributed voice and location representations as observed in the present study conceivably ensures sufficient flexibility to accommodate sound encoding in changing acoustic environments and with changing behavioral goals^44^.

Finally, we showed that sensitivity to a talker’s location features is similar across sites that are at lower stages of the hierarchy and sites that are at higher stages of the hierarchy. These results deviate from the view that spatial sensitivity emerges only in higher-order regions belonging to the functionally specialized location stream^32,40^. Instead, our findings are in agreement with more recent studies with active task designs which demonstrated that neural location sensitivity in early processing stages (i.e. primary auditory cortex) is more pronounced during active, goal-oriented localization^8,9^. Taken together, our results emphasize that experiment designs comprising active tasks and naturalistic stimuli are crucial to uncover representational mechanisms related to goal-oriented behavior in complex auditory scenes.

### Pre-attentive and attentive linking of voice and location to form complete auditory objects

Whether attention is required for auditory object formation remains a matter of debate^2,15^. Some have argued that auditory streams are formed pre-attentively, for example by the activation of separate populations of neurons^23^. Others have posited that attention is required to bind together the various attributes of the attended object^15^. Our data showed that a subset of cortical sites exhibited sensitivity to both voice and location features (‘dual-feature sensitive sites’), similar to prior findings of multi-feature sensitivity in ferret auditory cortex^13^. Moreover, we showed that the population response patterns of these dual-feature sensitive sites gave rise to the representation of the multi-dimensional auditory object, that is, the localized talker. To what extent this mechanism is pre-attentive requires further investigation.

In addition, we found that pairs of single-feature voice or location sensitive sites – corresponding to the ‘feature analysis’-stage of the temporal coherence framework^15,45,46^ – exhibited a build-up of temporal coherence over time both with and without attention to their preferred features. However, attention to the preferred features of the pairs of sites enhanced temporal coherence between these sites compared to the unattended condition. Temporal coherence started to increase 2 s after stimulus onset, in line with prior reports of perceptual and neural signatures of stream formation^37,38^. The observed increase of temporal coherence without attention, conceivably a result of synchronization to the temporal structure of the speech stimuli^47^, may facilitate pre-attentive stream formation. However, the boost in temporal coherence with attention to the preferred features conceivably reflects top-down attentional modulation of feature binding. This result is consistent with accumulating evidence^16,18^ supporting temporal coherence as a potential mechanism for grouping of perceptual features.

We posit that the observed increase in temporal coherence with attention to the preferred features emerges here through stronger phase alignment of the neuronal responses of voice- and location-sensitive sites to the stimulus’ temporal profile when attention is directed towards their preferred feature. Similar mechanisms have been reported previously for linking auditory and visual information^48,49^. As the present experimental set-up does not permit evaluation of this hypothesis (that is, the number of trial repetitions is not sufficient), future experiments with a higher number of stimulus repetitions are needed to accurately quantify phase alignment in this context and to assess the relationship of phase alignment to temporal coherence.

Taken together, our data indicate that linking of a talker’s voice and location features in spatial multi-talker scenes emerges from a mixture of (potentially pre-attentive) activation of dual-feature sensitive neural sites, population coding and attentional modulation of temporal coherence between voice and location sensitive sites.

### A continuum of attentional modulations of voice and location encoding

In agreement with prior work^21,25^, our results show that attending to a talker’s voice and location elicited weak attentional response gain control. Further, attention dynamically changed spectrotemporal tuning properties of late-response cortical sites, resulting in contrast matched filtering shape changes that enhanced local selective tracking of the attended talker’s speech. These results connect prior work in animals which showed that task performance and attention changed spectrotemporal tuning in auditory cortex^35,50^ to attended speech encoding in complex scenes in human auditory cortex. Moreover, these results extend findings from prior neural measurements in human auditory cortex, which showed that contextual information elicited adaptive STRF tuning to boost perception of degraded speech^51^.

Further, an attended talker’s location features elicited comparable attentional gain control in early- and late-response sites, suggesting that attention affects spatial processing at low-level as well as higher-order processing stages. More research is needed to establish whether these spatial attention effects in low-level cortical regions emerge from feedback projections originating in higher-order regions^44^. Additional work is also needed to evaluate whether attending a talker’s location in multi-talker scenes affects spatial tuning. That is, studies using single-source experiment designs with an active listening task reported sharpening of spatial tuning in primary auditory regions^8,9^ and it is likely that similar effects take place in multi-talker scenes to support segregating background from foreground. However, to assess this hypothesis, an experiment design with more fine-grained sampling of azimuth locations is required to elucidate attentional modulation of spatial receptive fields.

### Conclusion and outlook

Our results point to distributed and joint voice and location encoding across auditory cortex during active, goal-directed behavior. These findings support the view that object formation and attentional selection emerge gradually and in a distributed manner from the auditory hierarchy, rather than at one specific site or region in auditory cortex^3^. Such a distributed code flexibly accommodates rapid changes in the (acoustic) environment as well as changing behavioral goals. Crucially, the present findings demonstrate the need for real-life, complex stimuli and experimental designs including active behavioral tasks to understand cortical processing of multi-dimensional auditory objects. Future studies including stimuli spanning a larger and more fine-grained range of talkers, locations and other sound features can further unravel local cortical tuning properties as well as population representations of multi-dimensional auditory objects. Finally, complementing sEEG measurements with high-density intracranial measurements (e.g. high-density electrocorticography [ECoG], e.g. ^52^) are critical to refine cortical maps of local feature sensitivity, to tease apart fine-grained population representations within and across macro-anatomical regions, and to further our insights into feature binding through temporal coherence.

## METHODS

### Participants and data collection

We analyzed the data of seven adult neurosurgical participants which were implanted with high-density multi-electrode arrays (3.5 mm electrode spacing) as part of a clinical procedure to localize the focus of their epilepsy. As the data collection procedure has been described previously in detail^25^, we provide a brief summary here. Participants were fluent American English speakers and reported normal hearing. The Institutional Review Board of Feinstein Institute for Medical Research approved the research protocol. All participants signed an informed consent form prior to participation in the study. Three participants were implanted bilaterally, four participants were implanted in the right hemisphere. We recorded intracranial EEG signals at a sampling rate of 3,000 Hz, using either subdural or skull electrodes as reference. During recordings, speech stimuli were presented over headphones. Sound levels were adjusted for each participant to a comfortable level.

### Preprocessing

A detailed description of preprocessing of the neural data can be found in ^25^. In short, data preprocessing included montaging to a common average reference, noise removal, extraction of the high gamma envelope (70 – 150 Hz)^26,27^ using the Hilbert transform. Finally, neural responses were down sampled to 100 Hz and z-scored across single speaker blocks and across multi-talker blocks (i.e. calculated over both male and female trials, but separately for single- and multi-source blocks).

### Speech responsive electrodes

To assess which electrodes exhibited a robust response to speech streams, we computed for each electrode the mean baseline response as the average of the high-gamma envelope during 0.5 seconds preceding stimulus onset, and the mean speech onset response as the average of the high-gamma envelope in the 0.5 seconds following stimulus onset. To test for a statistically significant auditory response, we performed a paired samples t-test for each electrode and applied FDR correction across electrodes to correct for multiple comparisons. Only electrodes that exhibited a robust auditory response at *q* < 0.05 were included in the remainder of the analysis.

### Estimating spectrotemporal receptive fields (STRFs) and response latency

First, we computed a cortical spectrogram representation of each sound scene using a model of early cochlear processing and mid-brain auditory processing (NSL toolbox^53^). We modeled cochlear processing using a filter bank of 128 constant-Q filters that were spaced equally on a logarithmic axis ranging from center frequency (CF) = 270 Hz to CF = 7,246 Hz. Next, we modeled auditory midbrain processing by taking the derivative along the frequency axis, performing half-wave rectification and applying short-term temporal integration^53,54^. This approach accounted for the enhanced frequency selectivity as a consequence of lateral inhibition, as well as reduced phase locking, observed after midbrain processing. Cortical spectrograms were computed based on monaural stimulus waveforms (i.e. independent of sound location). The resulting spectrograms had a sampling frequency of 100 Hz and were down sampled to 50 channels to reduce the number of parameters.

We then estimated the spectrotemporal receptive field (STRF) by linearly mapping the cortical spectrogram to the evoked response using the STRFlab MATLAB Toolbox^55^ (http://strflab.berkeley.edu). For each electrode, we used the past 300 ms of a stimulus to predict the neural response at every time point using normalized reverse correlation. To prevent overfitting, we used a five-fold cross-validation procedure. We optimized sparsity and regularization parameters by maximizing the correlation between actual and predicted responses. Using the resulting STRFs, we defined the response latency for each electrode as the time point corresponding to the peak energy in the STRF.

### Decoding voice and location features in single-talker and multi-talker scenes

We trained a four-class classifier on population neural response patterns to jointly decode a talker’s voice and location features. The four classes corresponded to ‘male talker, left’, ‘male talker, right’, ‘female talker, left’ and ‘female talker, right’. We used frame-by-frame, regularized least-squares (RLS) classification^22,34^ which produced for each time frame a linear weighted sum of the population of neural responses for each class^22^. We trained and tested classifiers on the sustained responses only (i.e., excluding response onset effects from 0 to 500 ms). The class with the highest average classifier output over all frames in the trial was taken as the predicted class.

To decode a single talker’s voice and location features, we trained the classifier in a leave-two-trials-out cross-validation procedure on the single-talker data (corresponding to 25 folds). We computed classification accuracy as the average accuracy across the 25 folds. Further, to evaluate the statistical significance of classification accuracies, we performed a permutation analysis in which we randomly permuted the class labels and repeated the complete 25-fold cross-validation procedure. We iterated this process 2,000 times to create a null distribution of classification accuracy. Next, we tested whether the observed classification accuracy exceeds the 95^th^ percentile of the null distribution of permuted accuracies (one sided test). We computed *p* as the proportion of permuted accuracies that was equal to or larger than the observed accuracy.

To decode an attended talker’s voice and location features in spatial multi-talker scenes, we trained the four-class classifier on the multi-talker data using a similar procedure as described above. In multi-talker scenes, class labels consisted of ‘attended male talker, left’, ‘attended male talker, right’, attended female talker, left’ and ‘attended female talker, right’. We also assessed statistical significance using a permutation procedure similar to the permutation procedure for single talker data.

Finally, we calculated marginal accuracies for the voice and location feature dimensions by labelling accuracy based on a single feature dimension only, ignoring the other feature dimension. For example, to quantify the marginal accuracy for voice features, we calculated the percentage of the trials for which the correct voice class was predicted (i.e. female or male talker), ignoring the predicted location class (i.e. left or right). We computed the marginal accuracy also as the average across the 25 folds and used the permutation procedure described above to assess the statistical significance of the marginal accuracies.

### Attention-driven response gains in multi-talker scenes

For each electrode, we quantified the strength and direction of attentional modulations of cortical responses in the multi-source scenes evoked either by attending to a talker’s voice features or by attending to a talker’s location features using Cohen’s *d*. That is, similar to the quantification of single-talker feature sensitivity described above, we computed the mean response for each trial in the multi-source condition as the mean from 0.5 s post sound onset to 1.5 s post sound onset. Then, to test for attention-driven response gains for a talker’s voice features, we computed the effect size for the difference between the mean responses to all ‘attend male’ and ‘attend female’ trials, irrespective of the attended location of the trials (*n* = 50 each). To test for attention-driven response gains for a talker’s location features, we computed the effect size for the difference between the mean responses to all ‘attend left’ and ‘attend right’ trials, irrespective of the attended talker of the trials (*n* = 50 each).

### Quantifying temporal coherence

We assessed temporal coherence in slow fluctuations in stimulus evoked responses between pairs of voice sensitive and location sensitive sites. This analysis was performed on a within subject level. Five subjects contained multiple pairs of voice-location sites and were therefore included in the analysis. Because the high-gamma envelope is considered a signature of neural population responses^26,27^, we computed temporal coherence on the high-gamma envelope. Further, we quantified temporal coherence using the coherency coefficient, which is the mathematical equivalent in the frequency domain of the cross-correlation function in the time domain^38^. Specifically, the coherence coefficient is the normalized average cross-power spectral density between signals *x* and *y* across trials at frequency ω computed as^36^:

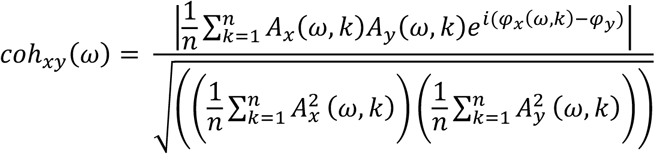

Here, we computed broadband temporal coherence over a frequency range of 2 – 22 Hz to map the development of attentional enhancement of temporal coherence over time, correspond to the range of slow fluctuations in which temporal coherence for feature binding is hypothesized to occur (i.e. 50 ms to 500 ms^15^). Furthermore, we computed narrowband temporal coherency for eight frequency bands with center frequencies 3, 6, 9, 12, 15, 18, and 21 Hz (bandwidth = 3 Hz) to examine the effect of attention on temporal coherence for specific frequencies.

## ACKNOWLEDGEMENTS

We thank Menoua Keshishian for sharing his expertise on STRF analysis. This study was supported by National Institute on Deafness and other Communication Disorders grant R01DC014279 (NM) and grant R01DC018805 (NM). This project has also received funding from the European Union’s Horizon 2020 Research and Innovation Program under the Marie Sklodowska-Curie grant agreement No 898134 (KH) and from the NWO Talent Program under the Veni grant agreement VI.Veni.202.184 (KH).

## SUPPLEMENTARY MATERIALS

### Supplementary Figure 1

Based on the observations of prior work^21,25^, we analyzed to what extent STRF properties of voice sensitive sites reflect tuning to the spectral profile of the preferred voice. Supplementary Figure 1 A shows that, as expected^21,25^, STRFs of sites responding preferentially to the male talker exhibited pronounced responses to distinct low frequency regions below 650 Hz, while sites responding preferentially to the female talker exhibited strong responses to frequencies between ∼280 and ∼1,400 Hz. Moreover, SRFs in Supplementary Figure 1 B show that spectral tuning properties of ‘male’-preferring sites exhibited an excitatory region at low frequencies between 50 Hz and 100 HZ, overlapping with F0 of the male talker (65 Hz). In contrast, the SRF of ‘female’-preferring sites exhibited an excitatory region between 160 Hz and 200 Hz, overlapping with F0 of the female talker (175 Hz). Thus, in line with previous reports^21,25^, we find that a voice site’s preference for either the male or female talker was driven by the correspondence between the spectral response profile of the site and the acoustic profile of the talker.

**Supplementary Figure 1.**
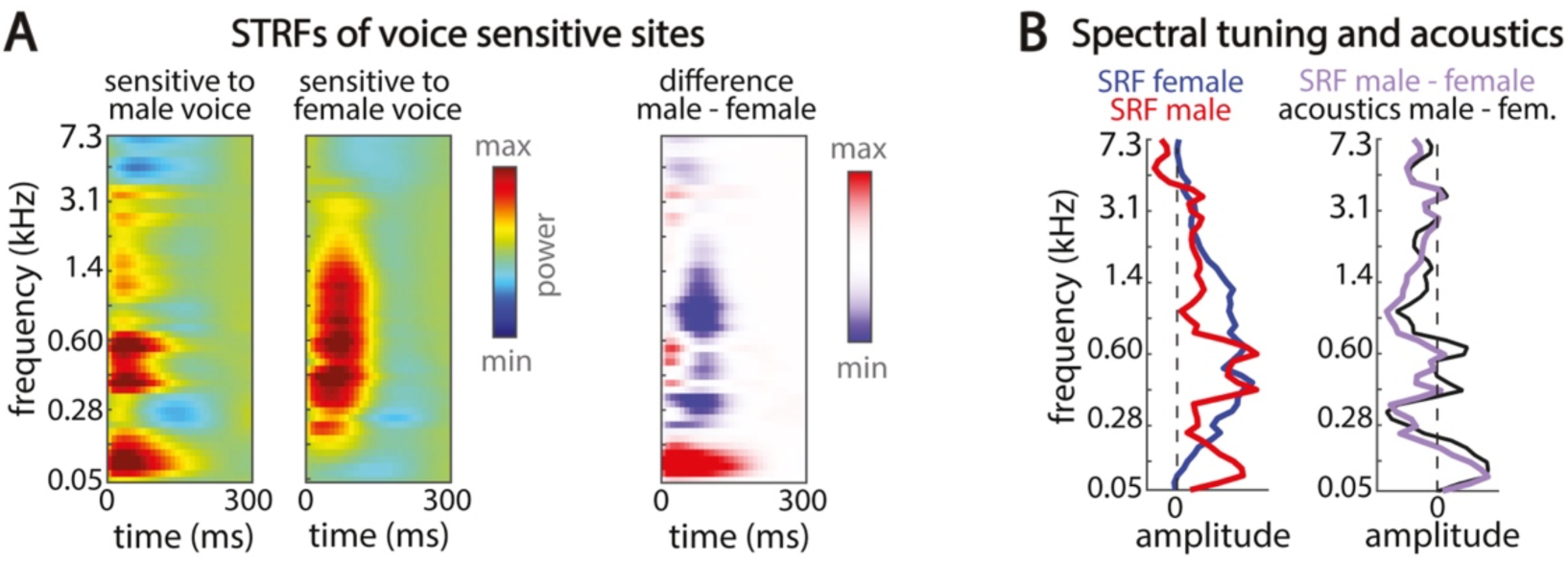
Spectrotemporal tuning characteristics explain sensitivity to a talker’s voice. (A) Spectrotemporal tuning properties related to voice sensitivity. From left to right: Average STRF for sites responding maximally to the male talker, average STRF for sites responding maximally to the female talker and the difference (STRF male – STRF female). (B) Comparing spectral tuning properties to the acoustics of the male and female talker. Left panel: Average spectral receptive field of sites responding maximally to a female talker (blue). Right panel: The correlation between the difference SRF (SRF male – SRF female) and the difference in the acoustics of the male and female talker.

### Supplementary Figure 2

**Supplementary Figure 2.**
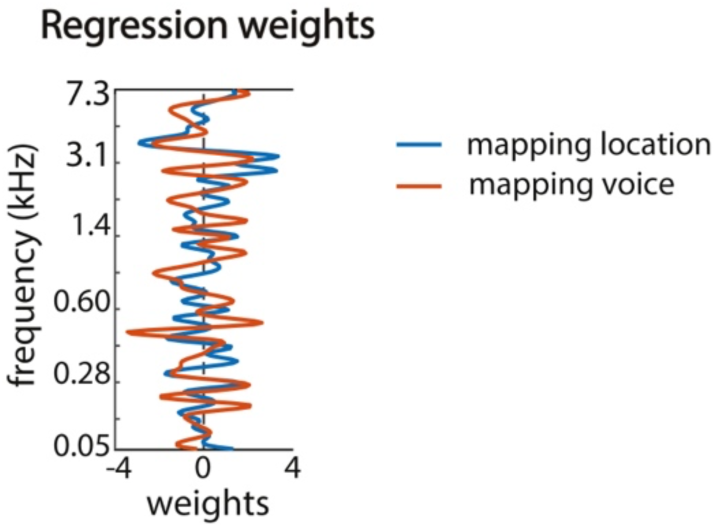
Regression weights applied to frequency bands to location and voice sensitivity from SRF. Blue line depicts regression weights across frequency bands learned to predict location sensitivity, orange line depicts regression weights learned to predict voice sensitivity. Weights showed a trend towards a weak correlation (*r* = 0.27, *p* = 0.06), indicating that most frequency bands had a different contribution to the prediction of location and voice sensitivity, while a few showed somewhat similar contributions to predicting voice and location sensitivity.

### Supplementary Table 1

**Supplementary Table 1.** The table below provides the distribution of speech responsive sites across subjects and cortical regions.

| Subject | Heschl’s gyrus (HG) | Planum temporale (PT) | Superior temporal gyrus (STG) |
| --- | --- | --- | --- |
| S1 | 4 | 6 | 4 |
| S2 | 11 | 5 | 18 |
| S3 | 2 | 2 | 11 |
| S4 | 3 | 1 | 19 |
| S5 | 4 | 10 | 13 |
| S6 | 7 | 8 | 5 |
| S7 | 7 | 3 | 4 |

## Notes

### Competing Interest Statement

The authors have declared no competing interest.

### Summary of Updates

- the analysis of spectrotemporal tuning properties - the analysis of temporal coherence - improvements in the text

